# Antibody maturation in germinal centers selects mutants based on BCR-antigen bond mechanical resistance

**DOI:** 10.64898/2026.03.11.710783

**Authors:** L. Awada, J. Sleiman, J.-M. Navarro, R. Torro, M. Biarnes-Pelicot, E. Hector, L. Limozin, C. Dong, P. Milpied, P. Robert

**Author notes:** Authors designed by an asterisk contributed equally (J. Sleiman, J.-M. Navarro, P. Milpied, P. Robert).

## Abstract

Antibody maturation in germinal centers (GCs) is traditionally viewed as a process that enhances antigen-binding affinity of B cell receptors (BCR). However, recent studies challenge this paradigm, revealing no systematic antibody affinity improvement during GC selection. Here, we investigate whether mechanical resistance of antibody-antigen bonds, rather than affinity, is the selective parameter driving GC maturation. Using biolayer interferometry measurements for assessing affinity and laminar flow chamber for mechanical resistance, we analyzed three lineages of mouse antibodies generated after immunization with ovalbumin. While affinity changes were heterogeneous, ranging from gains to losses across lineages, mechanical resistance consistently increased after maturation. Bond lifetimes under physiological forces (10-70 pN) converged to similar values across lineages, suggesting a selective pressure for mechanical stability. Functionally, NK cell activation in a surrogate *in vitro* ADCC assay correlated with bond lifetimes under force, but not with affinity. This indicates that lymphocytes are sensitive to the mechanical stability of antibody-antigen interactions, which may underlie GC selection. Our findings reconcile antibody maturation with the idea of enhanced binding, while aligning with the role of mechanical forces in B cell signaling and antigen uptake. These insights provide a framework for understanding GC biology and may inform the design of therapeutic antibodies.

## INTRODUCTION

Antibody maturation is a step of the adaptive immune response occurring in secondary lymphoid organs following initial B lymphocyte activation and proliferation triggered by the encounter of a specific ligand with its B Cell Receptor (BCR). A fraction of the resulting clones initiate the formation of germinal centers (GC) and start cycles of proliferation and mutation of the DNA coding for BCR sequences^1^, followed by a selection process which results in selected cells either re-initiating a cycle or exiting the GC after differentiating to antibody-secreting plasma cell (PC) or memory B cell. Only B lymphocytes able to uptake antigen can survive and enter the next cycle, as a necessary survival signal must be obtained by presenting peptides from the antigen to surrounding antigen-specific T follicular helper cells^2–5^. The capture and endocytosis of the antigen presented at the surface of follicular dendritic cells is thus a competitive step driving selection. The current paradigm established by Eisen^6^ is that several such cycles may select BCRs binding their antigen more efficiently, resulting in an improvement of affinity from the initial BCR of the lineage, and therefore called “affinity maturation”. However, observations from recent lineage tracing studies are incompatible with this hypothesis of affinity-driven antibody maturation, casting doubt on the idea of a Darwinian process that improves the binding properties of antibodies. El Tanbouly *et al* and Sprumont *et al* found no indication of affinity-based selection in the transition from GC B cell to PC; Sutton et al showed that the initiation of PC differentiation in the germinal center occurs in an affinity-independent manner ^7–9^.

Affinity, k_on_ and k_off_ quantify the binding of molecular species in solution, where molecular motions are in three dimensions (3D) and governed by diffusion due to thermal energy. Methods such as Surface Plasmon Resonance^10^ and BioLayer Interferometry^11^ (BLI) measure these parameters, that may accurately quantify the molecular interactions involved in the blocking function of antibodies. However, some steps of antibody and BCR interactions with their ligands do not occur in solution. During Antibody Dependent Cell Cytotoxicity (ADCC) by NK cells and Antibody Dependent Cell Phagocytosis (ADCP) by macrophages, the antibody and antigen are both linked each to a different cell or opsonized particle surface^12,13^. Also, BCR-antigen interactions during antigen presentation by a dendritic cell form while both molecular species are bound each to a different cell surface^14^. Interactions between two proteins linked each to a cell surface differ from interactions occurring in solution^15^. Bond formation occurs between molecules diffusing in two dimensions (2D) rather than three. Bond rupture is partly due to thermal energy, but also to the work of forces resulting from motions of the membranes, molecular motors and steric repulsion. A consequence is that affinity and kinetics constants measured in solution may not be relevant descriptions of binding properties in this context^16,17^. These are better described as functions of the mechanical force applied^18^ and need specific quantification methods such as Atomic Force Microscopy^19^, optical tweezers^20^ or laminar flow chambers^19,16,17,21^. During antibody maturation, follicular dendritic cells present at their surface the native antigen that B lymphocytes need to extract through their BCR to process it and present as peptide-MHC II complexes to T_FH_ cells. Mechanical forces are indeed involved, as BCR signaling depends on the application of forc^22,23^ and as B lymphocytes generate contractile forces to extract antigens^24,25^. Also, the stiffness of antigenpresenting cells influences antigen discrimination by BCR^25,26^, and the mechanics of the GC B cell immune synapse are specifically altered to increase force exerted on individual bonds^2,14^. Yet, to our knowledge, there are no studies about how antibody maturation in the germinal center impacts mechanical resistance of antibody-antigen bonds.

The present study aimed at testing the hypothesis that antibody maturation alters the mechanical properties of antibody-antigen bonds rather than their affinity, by quantifying and comparing the binding properties of antibodies at various steps of maturation both in solution and under mechanical force. In solution, we observed very diverse evolutions of affinity both in relative and absolute manners. By contrast, we observed a consistent gain in force resistance of antibody-antigen bonds defined as the half-life under a given force, that was reaching similar values between lineages, compatible with a force-driven selection process. On a functional level, force resistance correlated with the level of activation of NK cells triggered by ADCC on antigen-covered surfaces while affinity did not, suggesting that antibody maturation based on force resistance in the GC is important for immune effector functio..

## RESULTS

### Production of recombinant antibodies from BCR clonal lineages undergoing maturation in germinal centers

In order to study the antigen binding properties of BCR at different stages of maturation during GC reactions, we performed single-cell sequencing of RNA and BCR sequences for murine lymph node GC B cells responding to immunization with chicken ovalbumin (OVA) as a model antigen, as described in ^15^. These antibodies were produced in eukaryotic vector, as well as variants with reverted mutations representing the germline of each lineage. The antibodies were chosen based on their ability to bind ovalbumin assessed using flow cytometry and ELISA^15^. We chose 14 antibodies across 3 lineages and produced them as represented in Supplementary Figure S1, the sequences being given in Supplementary Table 1. The first lineage consisted of antibodies c179.p7.BC46, c179.p7.BC49, c179.p7.BC76, and c179.p7.BC75 sequenced from GC B cells harvested 10 days after a secondary immunization, and the inferred germline antibody c179. The second lineage consisted of antibodies c127.p9.BC18, c127.p9.BC41, c127.p9.BC50 sequenced from GC B cells harvested 10 days after a primary immunization and the inferred germline antibody c127. The third lineage consisted of antibodies c87.p8.BC19, c87.p8.BC27, c87.p8.BC50 and c87.p8.BC83 sequenced from GC B cells harvested 10 days after a secondary immunization and the inferred germline antibody c87. Antibodies from lineage c179 displayed between 15 to 19 mutations by comparison with their germline; among them 5 to 7 were in CDR regions (Supplementary Table 2). Antibodies from lineage c127 displayed between 3 to 9 mutations by comparison with their germline; among them one at best was in CDR regions. Antibodies from lineage c87 displayed between 10 to 17 mutations by comparison with their germline; among them, 3 to 4 were in CDR regions.

### Antibody maturation of three antibody lineages results in gains or losses in affinity with heterogeneous end-values

We measured affinity, association kinetics, and dissociation kinetics of those antibodies with the OVA antigen in solution using biolayer interferometry (Figure 1A, 1B, 1C and Supplementary Figure S2). The four matured antibodies of lineage c179 had affinities close to 10^−8^M, while the c179 germline antibody was not affine enough to trigger a signal with this method, maturation resulting in an obvious gain in affinity (Figure 1D). The three matured antibodies of lineage c127 had affinities in the order of 10^−6^M, c127 germline antibody also had an affinity in the order of 10^−6^M. Importantly, a significant loss in affinity was observed between germline and matured antibodies c127.p9.BC18 and c127.p9.BC50, while c127.p9.BC41 affinity was not significantly different from germline antibody affinity (Figure 1E). Two mature antibodies of lineage c87 had affinities in the order of 10^−6^M, showing a significant gain from germline antibody affinity, while c87.p8.BC19 had an affinity in the order of 10^−5^M that was not significantly different from germline antibody (Figure 1F). Dissociation kinetics in solution (off-rate or k_off_) measured in BLI showed similar results, accounting for most of the affinity changes; association kinetics showed mostly changes of smaller magnitude. In summary, when measured in solution, the relative outcome of antibody maturation within each lineage varied with either gain, loss, or little changes in affinity; the absolute outcome of antibody maturation also differed strongly between lineages, affinities of matured antibodies ranging from 10^−6^M to 10^−9^M.

**Figure 1:**
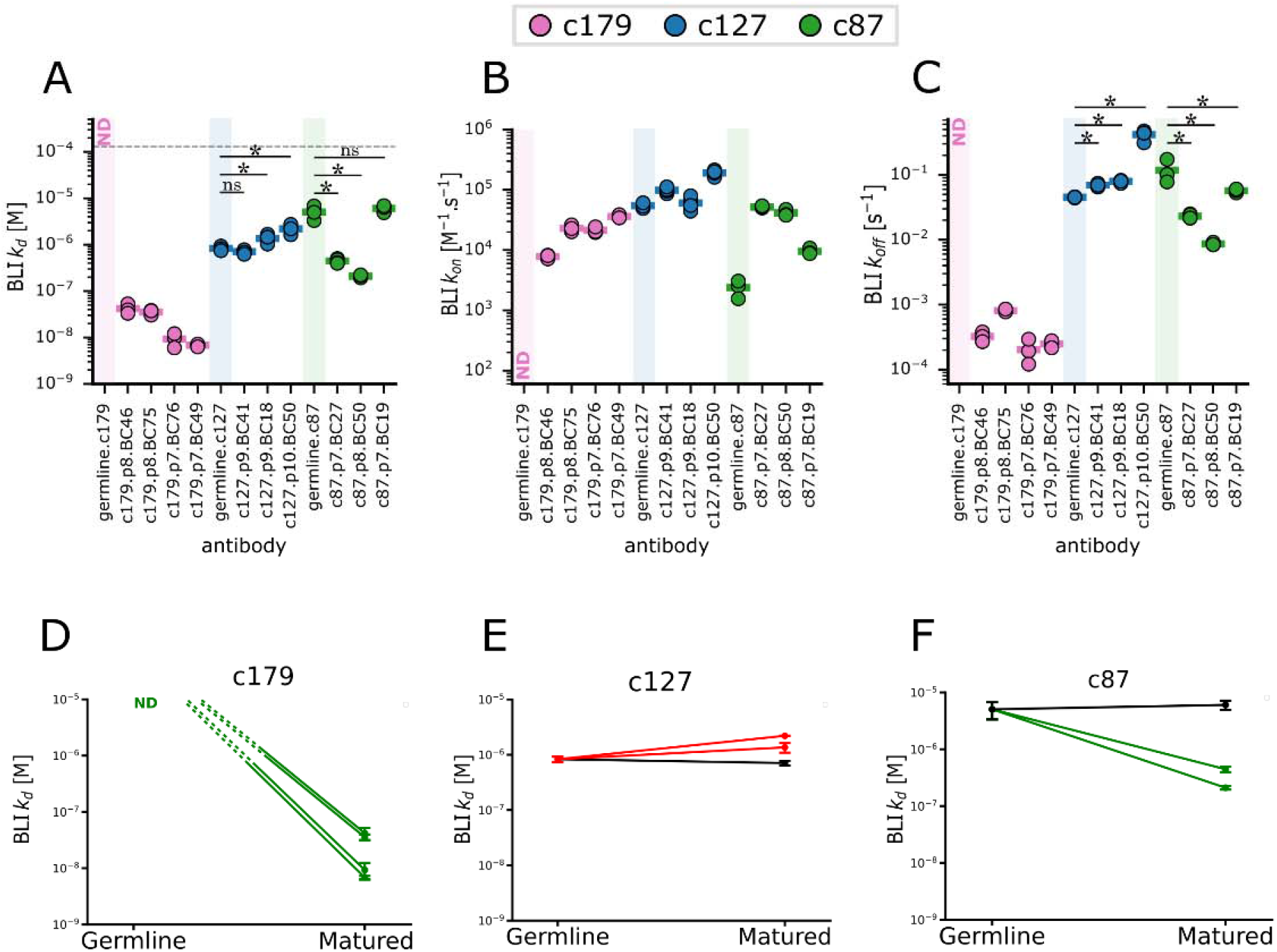
Measurement of antibody-antigen bonds in solution shows heterogenous evolutions of affinity, and heterogenous end-values of affinity after maturation. 3D binding parameters of antibody-antigen pairs across the c179, c127, and c87 lineages were measured by BLI. Each color corresponds to a different lineage; each point is an independent experiment, the horizontal line for each antibody corresponds to the mean of the distribution of the shown kinetic parameter. ND stands for non-detected. Statistical significances of differences were assessed using a Mann–Whitney test; *represents p-value <0.05. **(A)** shows the affinity constants K. **(B)** shows the on-rate (k_on_) values. **(C)** shows the off-rates (k_off_) values. **(D)**Comparison of affinity constants K of the single germline c179 antibody with the K values of its different mutated variants within the same lineage. Lines connect each germline-mutated antibody pair; green indicates a significant affinity gain and red indicates a significant affinity loss relatively to the germline, error bars are SD. **(E)**Similar data for all antibodies of the c127 lineage. **(F)**Similar data for all antibodies of the c87 lineage.

### Antibody maturation of three antibody lineages results in systematic gains in force resistance with similar end-values

We measured the binding properties of surface-bound antibodies to surface-bound OVA in single molecular bond conditions under forces, using our laminar flow chamber apparatus. In these experiments, antibody-covered microspheres were perfused under shear flow in chambers, moving after sedimentation over a surface bearing a low amount of OVA antigen (Figure 2A, Supplementary Figure S3). Binding of a single antibody to an OVA molecule would stop the microsphere, the duration of this arrest event being a direct measurement of the bond lifetime under the hydrodynamic force exerted by the shear flow on the microsphere. For each antibody of each lineage, we first determined the experimental range of OVA concentrations used during surface preparation resulting in single molecular bonds, as described in methods section (Figure 2B and Supplementary Figure S4). We measured dissociation kinetics (off-rate or k_off_) under five different forces (10pN, 25pN, 40pN, 55pN, 70pN) in single molecular bond conditions for each antibody (Figure 2C and Figure 2D). Under a 10pN force, germline antibodies of the three lineages showed dissociation significantly faster than matured antibodies. Matured antibodies had off-rates in the order of 0.1 s^−1^ irrespectively of the lineage. Germline antibodies bonds were no longer measurable beyond 10pN, and all matured antibodies were slip bonds, as their dissociation rate increased with force (Figures 2E and F). The relative outcome of antibody maturation was therefore very similar between lineages as maturated antibodies were significantly more force-resistant than germline antibodies. The absolute outcome of antibody maturation under force was also close between lineages, as off-rates of all matured antibodies of the three lineages were within the same order of magnitude.

**Figure 2:**
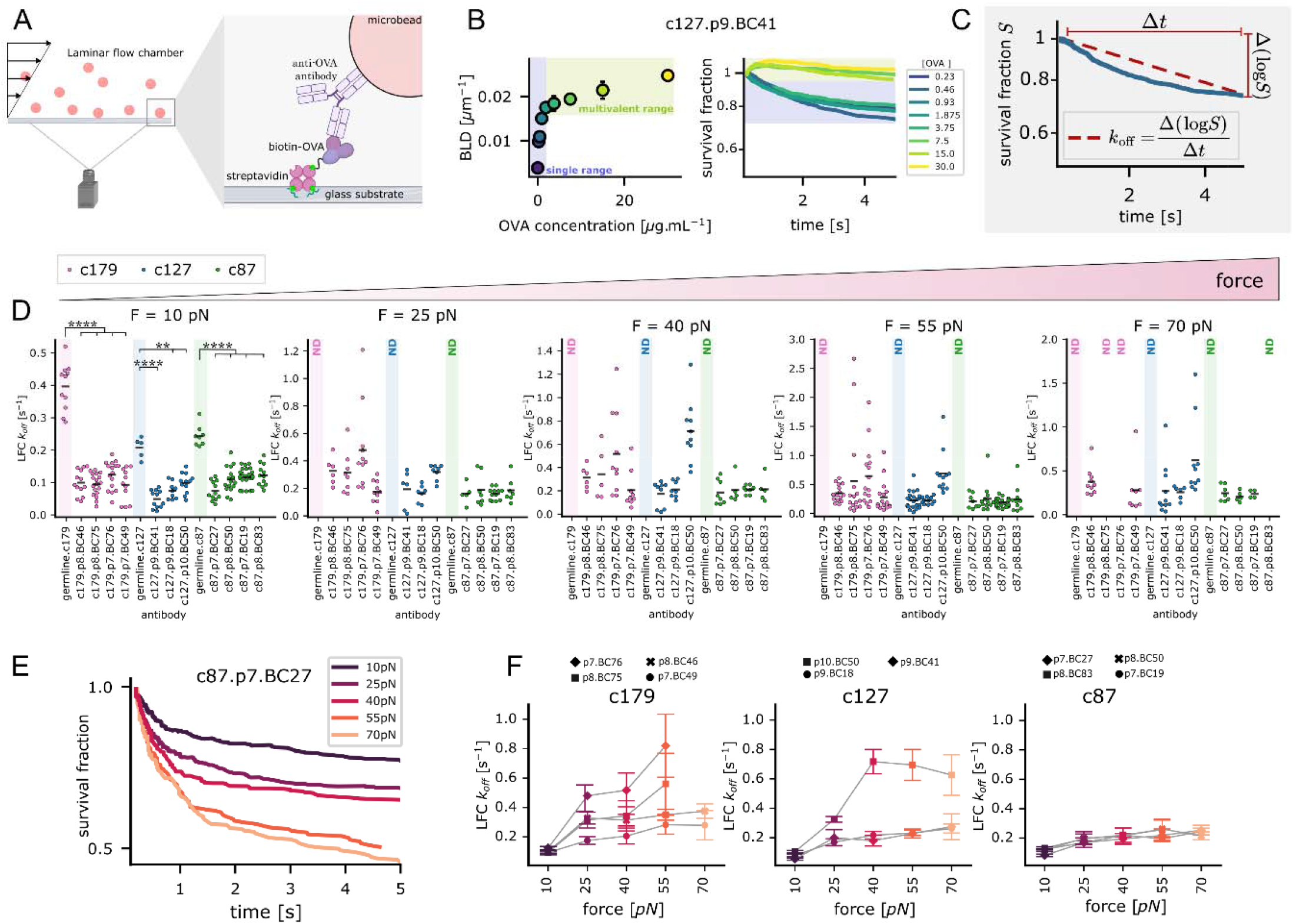
Measurement of antibody-antigen bonds under different forces reveals decrease in dissociation under force of the matured antibodies relative to their germline for each lineage, as well as similar end-values of dissociation under 10pN force. **(A)**Principle of the laminar flow chamber experiment. Antibody-coated microspheres move in a shear flow above a low-density ovalbumin-coated surface. A single antibody-ovalbumin bond may stop the microsphere, with shear flow exerting a force on the bond. **(B)** Assessment of single molecular binding range exemplified by c127.p9.BC41. Left, BLD values versus concentration of ovalbumin for 10pN force on bonds. Right, logarithm of the fraction of surviving bonds versus duration of the bonds under a 10pN force. **(C)** Calculation used to estimate the off-rate (k_off_) of each bond under a given force. **(D)** Off-rates (k_off_) measured across the 3 lineages, grouped by force. Each color corresponds to a lineage; each graph represents the off-rate (k_off_) distributions of the tested antibodies at a force range; each point is a replicate experiment. The horizontal line at each force corresponds to the mean of the distribution of off-rate (k_off_) values. Statistical significances of differences were assessed using Student’s t-test; ** represents p value <0.01 and **** represents p value < 0.0001. **(E)** Observation of slip bonds exemplified by c87.p7.BC27 antibody: survival fraction at a given time decreases when force increases. **(F)** Off-rates (k_off_) values measured under different forces for the antibodies of the c179, c127, and c87 lineages. Each point corresponds to the mean of the distribution of k_off_ values as measured in **C** under a given force; error bars are SD.

### Partially reversing mutations gained during antibody maturation reduces force resistance as well as affinity in a way compatible with a progressive selection process

Lineage c179 displayed the largest gain in affinity after maturation. We generated mutants of this lineage to produce putative variants intermediate between germline and matured antibodies (Figure 3A). The sequence of the Nearest Common Ancestor (NCA) was inferred from sequences of antibodies c179.p7.BC46, c179.p7.BC49, c179.p7.BC76, and c179.p7.BC75 sequenced from harvested GC B cells. From NCA sequence were also derived a mutant variant expressing the framework mutations of NCA but lacking all its CDR mutations (FW+/CDR-), and a mutant variant lacking the framework mutations of NCA but expressing all its CDR mutations (FW-/CDR+). Two glycine to aspartic acid mutations present in NCA heavy chain CDR1 and CDR2 were of possible importance as they each added a negative charge in the paratope. We produced a mutant (P1) deprived of both mutations, a mutant lacking the CDR1 mutation (P1.1), and a mutant lacking the CDR2 mutation (P1.2).

**Figure 3:**
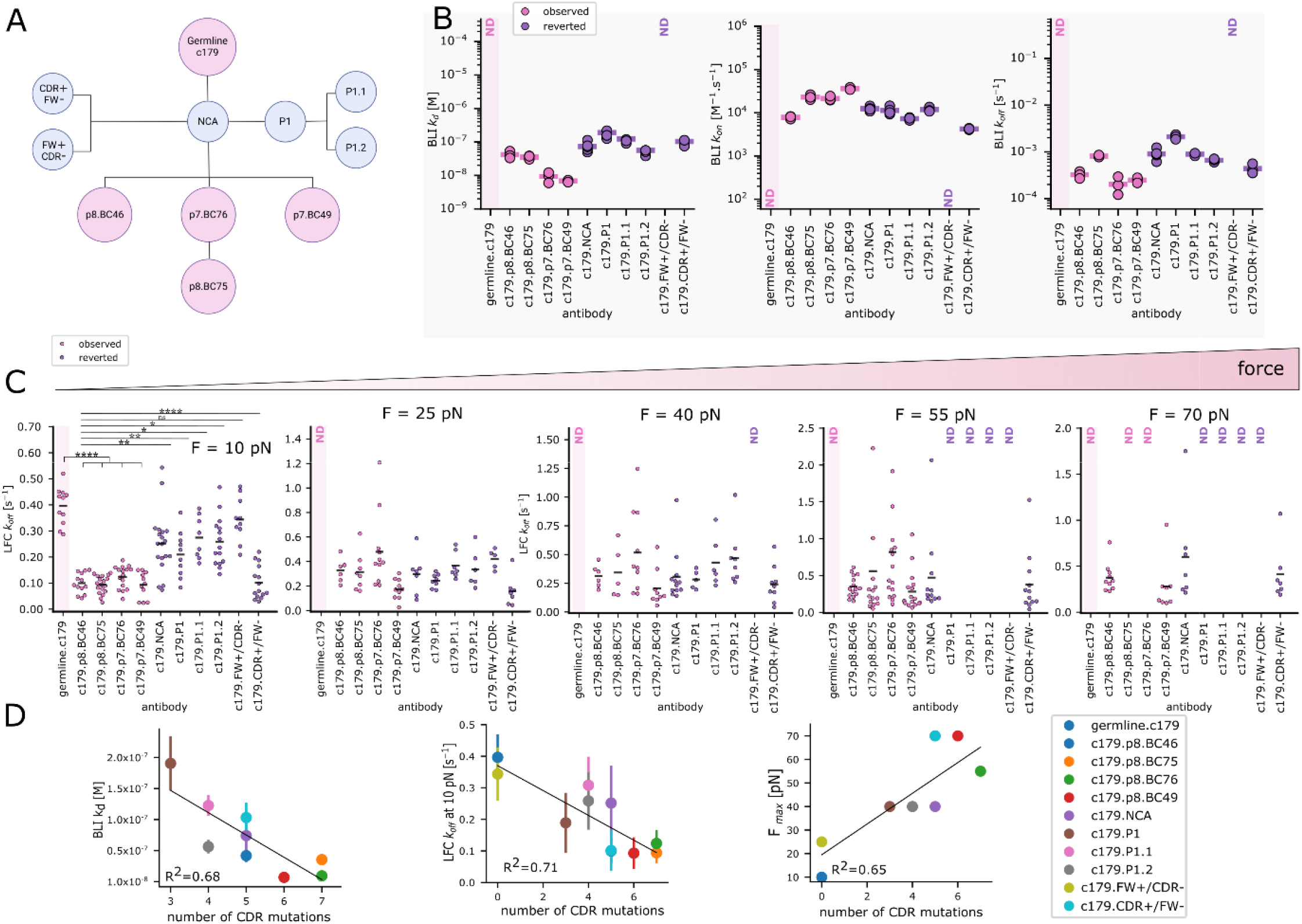
Antibody-antigen bonds with partially reverted mutations acquired during maturation have intermediate binding properties both under force and in solution. **(A)** Schematic phylogenetic tree of the c179 lineage with original and germline antibodies represented in pink and engineered antibodies expressing reverted mutations represented in purple. **(B)** Graphs of the affinity and kinetic parameters measured in solution for the observed antibodies and the antibodies with reverted mutations of the c179 lineage. Left graph shows K values, middle graph shows on-rate (k_on)_ values, and right graph shows offrate (k_off_) values. Each point represents an independent experiment. For each antibody, the horizontal line corresponds to the mean of the distribution of the shown kinetic parameter. Statistical significances of differences were assessed using a Mann–Whitney test. ND stands for non-detected. **(C)** K_off_ values of antibodies of the c179 lineage grouped by force. Each graph represents the off-rate (k_off_) distributions of the tested antibodies at a force range. The lighter pink shade represents the observed antibodies, and the darker purple shade represents the antibodies with reverted mutations. Each point represents an independent experiment. The horizontal line at each force corresponds to the mean of the distribution of off-rate (k_off_) values. Statistical significances of differences were assessed using Student’s t-test; * represents p-value <0.05, **represents p-value <0.01, and ****represents p-value <0.0001. ND stands for non-detected. (**D**) Relationship between the number of CDR mutations and binding properties measured across the c179 lineage. Each point corresponds to one mutated antibody variant compared to its germline ancestor. Left graph shows BLI K values as a function of CDR mutation number, middle graph shows LFC offrate (k_off_) at 10 pN as a function of CDR mutation number; error bars are SD in both cases. Right shows maximum force F_max_ allowing observation of single bonds as a function of CDR mutation number. Linear regressions and Spearman R^2^ determination coefficient values are shown for each dataset.

In solution (3D), we measured the binding of each of these six new antibody populations to ovalbumin using BLI (Figure 3B). NCA antibody and NCA-derived antibodies FW-/CDR+, P1, P1.1 and P1.2 showed intermediate affinity between germline c179 antibody (non-observable in BLI) and matured antibodies of c179 lineage. FW+/CDR- mutant binding to ovalbumin was not observable in BLI. We measured using the laminar flow chamber (Figure 3C and Supplementary Figure S5) the dissociation kinetics under forces ranging from 10pN to 70pN as in previous experiments. NCA antibody and NCA-derived antibodies FW-/CDR+, P1, P1.1 and P1.2 showed intermediate dissociation kinetics between germline c179 antibody and matured antibodies of c179 lineage. NCA-derived antibody FW+/CDR-showed no significantly different dissociation compared to the germline antibody. The number of mutations in CDR was broadly correlated to affinity, to off-rate under 10pN, and to the maximum force the bond could withstand (Figure 3D). Thus, antibodies with partially reversed mutations had both 3D and 2D binding properties intermediate between germline antibodies and matured antibodies, this being compatible with a progressive maturation process.

### NK cells activation by antibodies does not correlate with their affinity but correlates with their force resistance

To assess whether immune cells would be functionally sensitive to the differences in antibody binding properties measured in our experiments, we measured as a surrogate ADCC assay NK cell degranulation in response to ovalbumin surfaces coated with antibodies from two different lineages. We prepared sterile wells with bottom surfaces coated with biotinylated ovalbumin using a surface chemistry similar to laminar flow chamber coating, then saturated them with a solution of the chosen antibody. We harvested NK lymphocytes from mice spleens and sorted them by negative magnetic sorting, pre-incubated them with IL-2 overnight and deposited them in each well where they incubated for three hours. Cells would sediment on the surface, then eventually activate by interaction between NK lymphocyte CD16 receptors and Fc fragments of anti-ovalbumin antibodies. Expression of CD107a/LAMP1 was quantified by epifluorescence microscopy, allowing to determine a background threshold then to quantify the proportion of activated cells per antibody type (Figure 4A). We measured the proportion of activated NK cells (Figure 4B) for 6 antibodies of lineage c179, spanning the affinity range of the lineage and where maturation in lymph node resulted broadly in a large gain in affinity compared to germline, and for lineage c127 where maturation in lymph node resulted broadly in a small loss of affinity compared to germline. Neither affinity nor off-rate binding properties measured in solution correlated with NK cell activation (Figure 4C). By contrast, dissociation kinetics measured under a 10pN force correlated strongly with activation properties (R^2^ = 0.78, p = 0.0007, Figure 4D). Weaker correlation remained observable for dissociation kinetics measured under higher forces. In summary, NK cells in an ADCC-like assay were sensitive to the maturation-induced differences in force resistance, but not in affinity, that triggered differentially their functional response.

**Figure 4:**
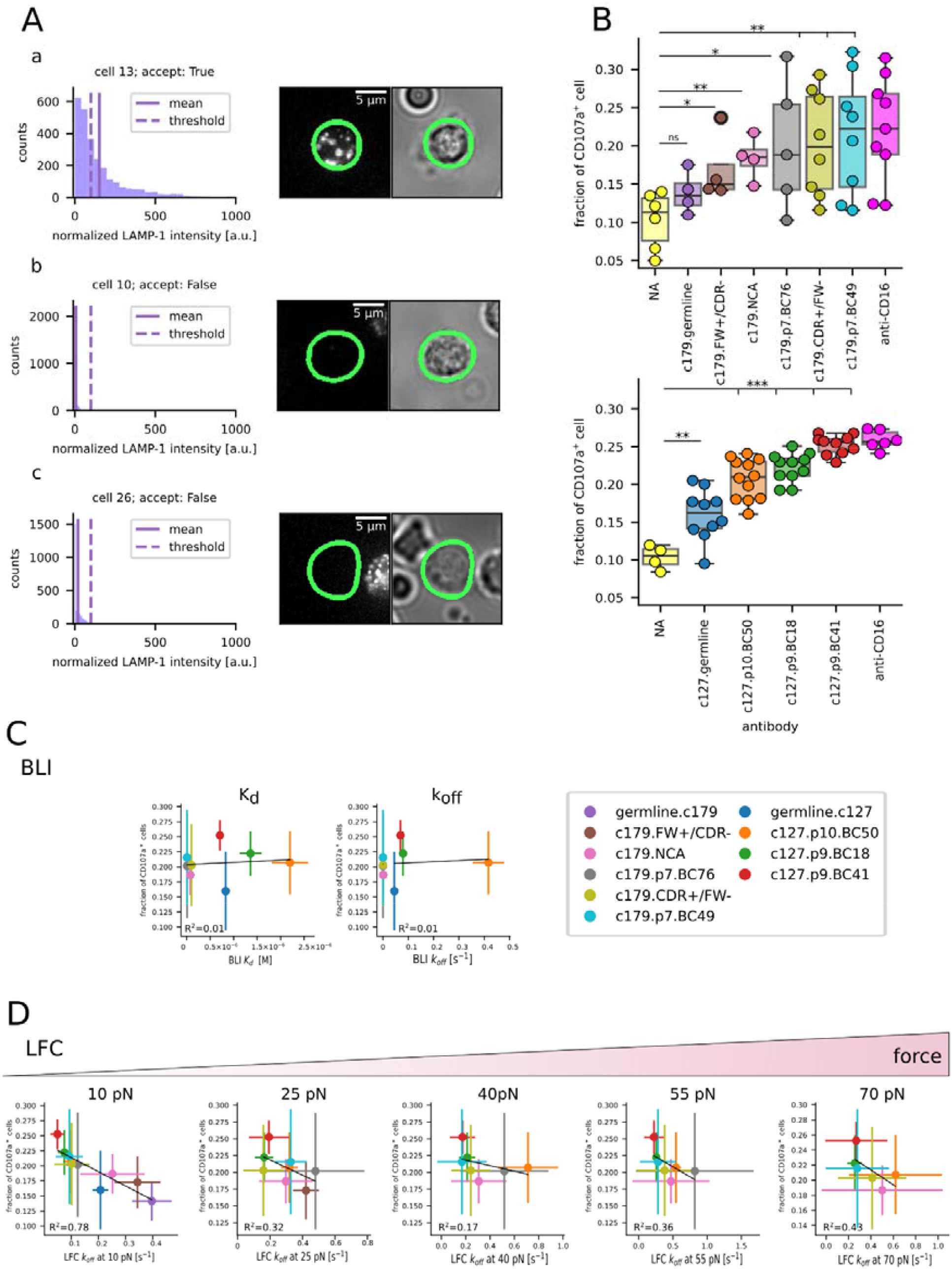
Quantification of NK cells activation in response to surfaces coated with ovalbumin and anti-ovalbumin antibodies. **(A)** Automated quantification of CD107a/LAMP1 detection. In **a**, on the left is the intensity histogram due to Alexa647 anti CD107a/LAMP1 antibody measured in epifluorescence in the area limited by a green contour in the leftmost photograph. The rightmost photograph is the transmission image allowing CellDetective to detect the cell contour, shown also in green. A threshold of mean intensity is defined so a mean 10% of cells in negative controls were considered expressing CD107a/LAMP1. Here, the cell signal overcame the non-specific binding threshold, and the cell is considered as expressing CD107a/LAMP1. In **b** and **c**, data are processed identically; cells are not expressing CD107a/LAMP1. **(B)** Proportion of cells considered as expressing CD107a/LAMP1 represented versus the nature of the surface, for c179 lineage on top and c127 lineage at bottom, with ovalbumin alone coated surfaces as negative controls and CD16 antibody coated surfaces as positive controls. Each circle represents an independent experiment in terms of surface preparation; cells gathered from a given mouse would be used for two of such experiments. Box plots show maximum data interval, SD and median for a given condition. **(C)** Correlations plots of fraction of CD107a/LAMP1-expressing cells versus off-rates measured under force for the experimental range of force. **(D)** Correlations plots of fraction of CD107a/LAMP1 expressing cells versus affinities (K_D_) measured in solution and off-rate (k_off_) measured in solution. For **C** and **D**, Spearman determination coefficients are shown; statistical significances of linear regressions were assessed using Student’s t-test.

## DISCUSSION

We compared the binding properties of matured antibodies produced from BCR sequences of GC B cells gathered in mice after immunization with ovalbumin to the binding properties of mutants designed to represent either the germline antibody or several putative intermediate steps of maturation, for a total of 20 different antibodies coming from three different B cell lineages. The effect of maturation measured in solution (in 3D) varied, as gain, loss and an absence of change in affinity were observed. In addition, the binding properties of matured antibodies were very heterogeneous between lineages, with differences of three orders of magnitude measured in our experiments. By contrast, binding properties measured under physiological forces (in 2D) showed a consistent reduction of antibodies off-rates after maturation within all three lineages, with partial reversal of maturation mutations resulting in a loss of mechanical strength. The absolute effect of maturation was also similar between different lineages, as matured antibodies from different germlines display similar off-rates under a given force. Such a homogeneous outcome is compatible with a selection process where dissociation kinetics under a 10 pN force would be the sensed parameter, rather than affinity. This result may reconcile the recent observations of very heterogeneous antibody affinities measured after antibody maturation^7,8^ with the idea of a Darwinian selection process in the GC linked to an enhancement of antibody binding properties. It is also well-aligned with the mechanisms of B cell selection in the GC, where mechanical extraction of antigen by B cells is required for antigen processing and presentation to cognate T_FH_^24,25,23,26^.

Several mechanisms may explain how a force-dependent binding parameter could drive cell selection during antibody maturation. B lymphocytes expressing a newly mutated BCR in a GC need to endocytose antigen to present antigen and ultimately survive. Endocytosis would exert a mechanical strain necessary for the unbinding of antigen from the follicular dendritic cell surface and require BCR-antigen bonds strong enough to withstand this forc^26^. A second hypothesis is that the B cell could actively probe antigens under force, to enhance discrimination between specific and non-specific antigens by providing additional information for ligand discriminatio^27,28^. In a third hypothesis, we observed that NK cell activation correlated best here with off-rate (or half-life) under a 10pN force of the antigen-antibody bond they probe. Several studies using different methods measured TCR-pMHC mechanical properties and observed that off-rate (or half-life) under a 10pN force correlates best with T lymphocyte response^21,29,30^. The force withstood by BCR during antigen recognition is also in the same order of magnitud^25^. We propose that withstanding a force in the order of magnitude of 10pN could be a necessary feature of cell surface-linked ligand-receptor bonds. Ligand-receptor bonds in solution must withstand thermal agitation to achieve biologically significant lifetimes and thus display Gibbs energies in the order of several k_B_T. Similarly, cell membrane anchored ligand-receptor bonds would need to withstand non-specific mechanical forces (arising from repulsion by glycocalyx and membrane fluctuations) to achieve biologically significant lifetimes. This force resistance could be a recurrent selective factor for membrane anchored ligand-receptor bonds; its magnitude would be in the order of 10pN for lymphocytes. Strikingly, the steady-state force exerted by glycocalyx repulsion on a single TCR-pMHC bond at the APC-T lymphocyte interface was estimated to be around 18pN in a theoretical work by van der Merwe and colleague^31^. Also, our results suggest that lymphocytes can detect the differences in bond lifetimes measured here under a 10pN force. This observation suggests a possible link between antibody maturation and the antibody functions occurring at cell-surface interfaces through Fc receptors, as a force-based selection process would ensure the ability for antibodies to trigger NK cells or phagocytes if force resistance requirement is the same.

### Limitations of the work

This work is limited to the study of three B lymphocyte lineages following immunization with a single antigen and may not represent a general case. Moreover, this antigen is a small protein that may properly represent the antibody response to small soluble molecules such as toxins, but maybe not what happens when B lymphocytes encounter whole viruses and bacteria on antigen-presenting cells. However, our results show that it is possible to reconcile the idea of a selection process in the GC during antibody maturation with an enhancement of the binding properties of antibodies resulting from this maturation. As shown in other systems, affinity may be irrelevant when quantifying binding under force; here dissociation kinetics under force seem to be both a physical parameter compatible with being the subject of a selection in the lymph node, and a good predictor of effector cell response.

## MATERIALS AND METHODS

### Anti-OVA antibody production as described in Ado et al

IgG1 monoclonal antibodies were produced using the protocol described by Ado and colleagues^15^. Briefly, Aicda-CRE-ERT2+/− × Rosa26-eYFP-lox-stop-lox+/− mice were immunized with the model antigen chicken ovalbumin (OVA) to investigate germinal center B cell responses at day 10 following primary or secondary immunization. Starting with a cell suspension, single GC B cells undergoing affinity maturation were sorted by Fluorescence Activated Cell Sorting (FACS) into 96-well plates for reverse transcription, cDNA barcoding, and amplification. A fluorescent labelled ovalbumin was used among the flow cytometry markers to allow an estimation of BCR affinity for sorted cells.

A portion of the single-cell cDNA of selected germinal centers cells was used to prepare 5’-end RNA-seq libraries that were sequenced using the FB5P-seq method (FACS-Based 5-Prime End Single-Cell RNA-sequencing) allowing the parallel analysis of transcriptome-wide gene expression and OVA-specific BCR sequences^32^. The cDNA of selected cells of interest was used as input for cloning heavy and light chain variable regions into mouse IgG1 antibody expression plasmid vectors. The corresponding antibodies were produced by transient transfection of Expi293™ eukaryotic cells followed by purification for subsequent functional assays.

Functional assays of each antibody production were performed by ELISA and are also detailed in detail by Ado and colleagues^15^. ELISA assays consisted in preparing ovalbumin-coated 96 wells plates, followed by rinsing, blocking with a PBS-BSA solution, incubation with serial dilutions of the tested antibody, rinsing and revelation using an HRP-conjugated anti-mouse IgG antibody and a TMB solution.

The 3 germline antibodies corresponding to each lineage and the 6 variants of c179 lineage with reverted mutations were cloned into the same plasmid antibody expression vector used to produce the mutated antibodies. The variable regions of the heavy and light chains were synthesized by Twist Bioscience (USA) and subsequently cloned into the respective vectors. Following purification, the germline antibodies underwent the same production process and functional assays as the mutated antibodies.

### Biolayer interferometry experiments

Biolayer interferometry assays were performed on the Octet R2 instrument (Sartorius) at 25°C with shaking at 1000 r.p.m, using Octet® Anti-Mouse Fc Capture (AMC) biosensors. The biosensors were provided as dry tips that were hydrated in the kinetic buffer (PBS: bovine serum albumin (BSA) 0.1%: Tween 0.02%) for at least 10 minutes before the start of any given experiment. The final volume for all the solutions was 200 µL/well. Assays were performed at 25 °C in solid black 96-well plates (Geiger Bio-One) with agitation set to 1000 rpm to minimize nonspecific interactions. Briefly, a given sensor was equilibrated to establish a first baseline, then 5 µg/ml of antibody in 1× kinetic buffer was used to load the antibody on the surface of the biosensors. A biosensor washing step was applied before the analysis of OVA association to establish a second baseline. Association kinetics of the interaction were measured while dipping the sensor in an OVA solution in kinetic buffer, then dissociation kinetics of the interaction was measured in the kinetic buffer. For each antibody, sensorgrams represented the shift in wavelength of the spectrum reflected by the sensor tip versus time, with association starting at the instant of contact of tip with ligand solution and resulting in an increasing signal, and dissociation starting at the following contact of tip with solvent solution alone and resulting in a decreasing signal. BLI sensorgrams are shown in Supplementary Figure S2 for the c179, c127, and c87 clones, and Supplementary Figure S5 for c179 clones expressed with reverted mutations. In each graphic, replicate experiments at a given concentration are denoted by different colored traces forming a set of broadly superimposing curves. A minimum of three replicate experiments was performed per antibody. Each experiment was conducted over three or four concentrations that spanned the binding regime (the level of signal increasing with the concentrations) forming a total of three or four sets of superimposing curves per graph. Kinetic responses were fitted to the 1:1 Langmuir kinetic model to obtain values for association (k_on_), dissociation (k_off_), and the equilibrium dissociation constant (K_d_). Dissociation wells were used only once to ensure buffer potency. Each sensor was regenerated up to 4 times using a 10 mM glycine solution with a pH of 2.3 to acquire replicate data at different OVA concentrations.

### Laminar flow chamber experiments

Sample preparation was done as follows: glass slides were rinsed twice in absolute ethanol then washed in a « piranha » solution composed of 70% H_2_SO_4_, 15% water, and 15% H_2_O_2_ for 10 minutes, then rinsed with 5 liters of deionized water. Glass slides were coated with poly-L-lysin (150-300kDa, Sigma Aldrich) at 100µg/ml in a 10^−2^ M phosphate solution, pH 7.4 for 10 minutes, then rinsed in PBS, then incubated with glutaraldehyde (2.5% in pH 9.5 0.1M borate solution, Sigma Aldrich) for 10 minutes, then rinsed in PBS, then incubated in a 100µg/ml biotinylated BSA solution in PBS (Sigma Aldrich), for 30 minutes, then rinsed in PBS, then incubated in a 10µg/ml streptavidin solution in PBS (Sigma Aldrich), then rinsed in PBS. Glass slides were then mounted in a home-made multi-chamber, thermoregulated, flow chamber device, forming nine independent chambers of 12mm length and 2×0.250mm^2^ section. Each chamber was then filled with a biotinylated ovalbumin solution in PBS with 0.1% BSA at a chosen concentration, using cascade dilution to deposit in our 9-chamber apparatus 8 different amounts of biotinylated ovalbumin in each chamber plus one chamber as a negative control. Microspheres (Dynal M-450 tosylactivated, ThermoFisher) were rinsed three times in a 0.1M borate solution, then incubated overnight in a 200µg/ml solution of chosen mouse antibody under constant agitation.

Flow chamber experiments were performed using our home-made automaton controlled by an Arduino Mega 2560 card (Arduino). The multi-chamber device was thermoregulated at 37°C by water circulation, set on an inverted microscope with a 10× lens (Leica) with a digital CCD camera (UEye, IDS), and chamber entry was connected to the piping and syringes assembly. For each independent chamber, the automaton performed seven cycles, each forming an experiment under a given shear flow. Cycles consisted of agitation of microsphere suspension, injection of microsphere suspension in the chamber, and, for 90 seconds, simultaneous movie recording at 50 frames per second with M-JPEG on-the-fly compression and shear flow set at a given shear. The shear value was automatically modified for each new cycle until all chosen shear conditions had been recorded. The chamber was manually disconnected, and the next chamber was connected; then the automaton was re-launched. Raw data were in the form of movies of microsphere motion. Each different antibody-ovalbumin couple bond was measured in 8 to 12 independent experiments, with measurement performed for 8 different deposited amounts of ovalbumin plus a negative control for each independent experiment.

Statistics of molecular bond formation and rupture were determined by counting the frequency and duration of microspheres arrests using home-written Java plug-ins incorporated in ImageJ (NIH, USA). Briefly, a microsphere was considered to be arrested if the position of its center of mass did not change by more than 1 pixel during 180msec, and if its velocity before the arrest was within the velocity range of a sedimented microsphere moving in the shear flow. An arrest was considered to continue as long as the arrest criterion was satisfied, which yielded a duration. The durations of arrests were pooled for experiments sharing identical antibodies on microspheres, identical amount of ovalbumin on the surface, and identical shear rate, to build survival curves of the arrests. The survival curves were built by counting the fraction of arrests exceeding the minimal duration versus time. The Binding Linear Densities (BLD) were defined as the number of arrests divided by the total distance travelled by the microspheres in the velocity range of sedimented microspheres. The BLD of specific association under a given condition (i.e., a given shear rate and a given ligand surface density) was calculated by subtracting from the BLD measured with assay surfaces the BLD measured with control surfaces (without ovalbumin). Specific survival curves were calculated by subtracting, for each time step, the corresponding survival fraction of non-specific arrests measured in control experiments, times the ratio of control BLD over assay BLD for the condition. Force applied on the bond depended on shear flow that produced both an hydrodynamic drag *R*, and a torque *Γ* on the microsphere. A lever effect makes the actual force F applied on the bond dependent on the length of the molecular bond L. Force on bond was calculated as 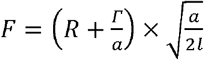 with *R* = 1.7005 × 6*π*μ*a*^2^*G* and *Γ* = 0.9440 × *π*μ*a*^3^*G* (with *R* the traction on the microsphere, *Γ* the torque on the microsphere, *a* the microsphere radius, *l* the total bond length (24nm), μ the medium viscosity (7×10^−4^ Pa.s at 37°C), and the shear rate^29,30^ (Supplementary Figure S3). Single molecular bond observation was assessed using arguments previously detailed ^30,33,34^: in a certain range of deposited amounts of ovalbumin, linear binding density was proportional to the amount of deposited ligand; survival curves of bonds would not change in this same range, demonstrating observation of similar binding events (Figure 2B, Supplementary Figure S4). Off-rates were calculated in a simple but robust way, as the opposite of the slope of the natural logarithm of each detachment curve, measured between 0 and 5 seconds (Figure 2C).

### NK cell activation assays

Murine NK cells were isolated from C57BL/6 (B6) mice spleens using a mouse NK isolation kit (Miltenyi Biotec, 130-115-818) according to the manufacturer’s protocol. Freshly isolated cells were subsequently cultivated overnight in Roswell Park Memorial Institute Medium (RPMI; Gibco by Thermo Fisher Scientific, Waltham, MA) 1640 supplemented with 25 mM GlutaMax (Gibco by Thermo Fisher Scientific, Waltham, MA),10% fetal calf serum (Gibco by Thermo Fisher Scientific, Waltham, MA), 100 U/mL penicillin (Gibco by Thermo Fisher Scientific, Waltham, MA), 100 µg/ml streptomycin sulfate (Gibco by Thermo Fisher Scientific, Waltham, MA), 1 mM sodium pyruvate (Gibco by Thermo Fisher Scientific, Waltham, MA), 50 µM *β*-mercaptoethanol (Gibco by Thermo Fisher Scientific, Waltham, MA), 10 mM nonessential amino acids (Gibco by Thermo Fisher Scientific, Waltham, MA), and 400 U/mL mouse IL-2 IS (Miltenyi Biotec, 130-120-662) in a 37°C incubator with 5% CO2. All cells were used the day following isolation and IL-2 overnight stimulation.

Volume-reducing inserts cut in 250µm thick PDMS sheet were deposited in an uncoated eight-well chambered polymer coverslip-bottom from Ibidi (80821, Ibidi) to form wells of 3 mm in diameter. Wells were initially rinsed with PBS, then incubated in a solution of biotin-labeled BSA (A8549, Merck) diluted in PBS at 100 µg/mL for 30 min at room temperature (RT) with agitation. Wells were rinsed four times with PBS, then incubated in a 10 µg/mL neutravidin solution (31000, Thermo Scientific) diluted in PBS for 30 min at room temperature with agitation. After a four-time PBS wash, the wells were coated with our biotinylated OVA diluted in 0.2% BSA at 10 µg/mL for 30 minutes at RT with agitation, followed by another four-time PBS wash. Subsequently, the wells were coated with 10µg/mL of the anti-OVA antibodies. For positive control experiments, surfaces were coated by biotin-labeled BSA and streptavidin as previously described, then incubated with 10µg/mL of biotinylated anti-mouse CD16 antibody (BioLegend). Finally, a suspension of 18000 mouse NK cells with anti-mouse CD107a/LAMP1 AF-647 at 10µg/ml (Biolegend, USA) was deposited in each well for 3 hours in an Okolab chamber where temperature was set at 37°C and CO2 level at 5%, before starting image aquisition.

Images were acquired using an AxioObserver (Zeiss, Germany) inverted microscope equipped with a Zeiss Plan-APOCHROMAT 40x/ 1.3 Oil Dic (UV) VIS-IR 420762-9800 lens, an Okolab chamber where temperature was set at 37°C and CO2 level at 5% and controlled by the Micromanager software (NIH, USA). For a period of two hours, sequences of 13 to 20 non-overlapping positions were repeatedly acquired per well. Each position was obtained at various focal planes (Z-planes) by increments of 2µm using the red fluorescence cube followed by a singular brightfield image, before transitioning to the next position and, subsequently, the next well.

Maximum intensity projections were generated from the z-stacks to consolidate the fluorescence signals. A custom Python script was used to correct background fluorescence. Cells were roughly segmented using a Gaussian blur (kernel: 4) and standard deviation (STD) filter (kernel: 4). A paraboloid model was then fitted to the non-cellular pixels using the package LMFIT and subtracted from the image. For model training, cells of interest were manually labeled in brightfield images using Labkit (Fiji plugin). These labeled images were used to train a custom StarDist model via a Jupyter Notebook. The trained StarDist model was imported into Celldetective software for segmentation and intensity measurement of CD107a/LAMP1 in all conditions^35^. Downstream analysis of the intensity data was performed using custom Python scripts in Jupyter Notebooks.

## ACKNOWLEDGEMENTS

The authors warmly thank Dr Patrick Chames for access to his biolayer interferometry apparatus, and Drs Kaitelyn Spillane, Mauro Gaya and Olivier Theodoly for most fruitful discussions. We are grateful to all past and present members of the “Integrative B cell Immunology” lab at CIML for useful discussions. This work was supported by grants from ANR (ANR-17-CE15-0009-01 “MoDEx-GC” and ANR-23-CE15-0025-01 “GCselection”) to P.M. This work was supported by institutional grants from INSERM, CNRS and Aix Marseille University to the CIML. LA and JS were supported by doctoral fellowships from the Turing Centre for Living Systems (CENTURI). EH was supported by a doctoral fellowship from the French Ministry of Research and Higher Education.

## AUTHORS CONTRIBUTIONS

LA and JS performed LFC and BLI measurements; LA set up, performed and analyzed cellular measurements with the help of MP. JMN produced antibodies and performed quality controls with the help of LA and JS. CD performed single-cell BCR sequence data preprocessing. EH performed BCR sequence analysis. RT and LL designed the CellDetective software. PR made the LFC apparatus and software suite; LA, JS and PR did LFC data analysis. PM and PR designed the study, supervised the study, acquired funding and wrote the manuscript. All authors revised the manuscript.

