## Supplementary information for "Antibody maturation in germinal centers selects mutants based on BCR-antigen bond mechanical resistance"

### Supplementary Table 1 :

| **Lineage c127 (IGHV1-64*01_IGKV1-99*01)** | |
|---|---|
| **Heavy chain sequence** | |
| germline | caggtccaactgcagcagcctggggctgagctggtaaagcctggggcttcagtgaagtt-<br>gtcctgcaaggcttctggctacactttcaccagctactggatgcactgggtgaagcagaggcctggacaagg<br>ccttgagtggattggaatgattcatcctaatagtggtagtactaactacaatga-<br>gaagttcaagagcaaggccacactgactgtagacaaatcctccagcacagcctacatgcaactcagcag<br>cctgacatctgaggactctgcggtctattactgtgcaagatccctc-<br>tactattccagaaggtactggtacttcgatgtctggggcacagggaccacggtcaccgtctcctca |
| p9.BC41 | caggtgcaactgaagcagtctggggctgagctggtaaagcctggggcttcaatgaagtt-<br>gtcctgcaaggcttctggctacactttcaccaactactggatgcactgggtgaagcagaggcctggacaagg<br>ccttgagtggattggaatgattcatcctaatagtggtagtactaactacaatga-<br>gaagttcaagagcaaggccacactgactgtagacaaatcctccagcacagcctacatgcaactcagcag<br>cctgacatctgaggactctgcggtctattactgtgcaagatccctc-<br>tactattccagaaggtactggtacttcgatgtctggggcacaggaacctcagtcaccgtctcgagc |
| p9.BC18 | caggtgcaactgaagcagcctggggctgagctggtaaagcctggggcttcagtgaagtt-<br>gtcctgcaaggcttctggctacactttcaccagctactggatgcactgggtgaagcagaggcctggacaagg<br>ccttgagtggattggaatgattcatcctaatagtggtagtactaactacaatga-<br>gaagttcaagagcaaggccacactgactgtagacaaatcctccagcacagcctacatgcaactcagcag<br>cctgacatctgaggactctgcggtctattactgtgcaagatccctc-<br>tactattccagaaggtactggtacttcgatgtctggggcacagggactctggtcactgtctcgagc |
| p10.BC50 | caggtccaactccagcagcctggggctgagctggtaaagcctggggcttcagtgaagtt-<br>gtcctgcaaggcttctggctacactttcaccagctactggatgcactgggtgaagcagaggcctggacaagg<br>ccttgagtggattggattgattcatcctaatagtggtagtgctaactacaatga-<br>gaagttcaagagcaaggccacactgactgtagacaaatcctccagcactgcctacatgcaactcagcagc<br>ctgacatctgaggactctgcggtctattactgtgcaagatccctc-<br>tactattccagaaggtactggtacttcgatgtctggggcacagggactctggtcactgtctcgagc |
| **Light chain sequence** | |
| germline | gatgttgttctgacccaaactccactctctctgcctgtcaatattggagatcaagcctc-<br>tatctcttgcaagtctactaagagtcttctgaatagtgatggattcacttatttggactggtacctgcagaagcc<br>aggccagtctccacagctcctaatatatttggtttctaatcgattttctggagttccaga-<br>caggttcagtggcagtgggtcaggaacagatttcacactcaagatcagcagagtggaggctgaggatttggg<br>agtttattattgcttccagagtaactatcttcctctcacgttcggtgctgggaccaagctggagctgaaa |
| p9.BC41 | gaaattgtgctcacccagtctccactctctctgcctgtcaatattggagatcaagcctc-<br>tatctcttgcaagtctcctaagagtcttctgaatagtgatggattcacttatttggactggtacctgcagaagcc<br>aggccagtctccacaactcctaatatatttggtttctaatcgattttctggagttccaga-<br>caggttcagtggcagtgggtcaggaacagatttcacactcaagatcagcagagtggaggctgaggatttggg<br>agtttattattgcttccagagtaactatcttccgctcacgttcggtgctgggaccaagctggagctgaaa |
| p9.BC18 | gatgttgtgatgacccaaactccactctctctgcctgtcaatattggagatcaagcctc-<br>tatctcttgcaagtctactaagagtcttctgaatagtgatggattcacttatttggactggtacctgcagaagcc<br>aggccagtctccacagctcctaatatatttggtttctaatcgattttctggagttccaga-<br>caggttcagtggcagtgggtcaggaacagatttcacactcaagatcagcagagtggaggctgaggatttggg<br>agtttattattgcttccagagtaactatcttccgctcacgttcggtgctgggaccaagctggagctgaaa |

| | |
|---|---|
| p10.BC50 | gatattgtgatgacccaaactccactctctctgcctgtcaatattggagatcaagcctc-<br>tatctcttgcaagtctactaagagtcttctgaatagtgatggattcacttatttggactggtacctgcagaagcc<br>aggccagtctccacagctcctaatatatttggtttctaatcgattttctggagttccaga-<br>caggttcagtggcagtgggtcaggaacagatttcacactcaagatcagcagagtggaggctgaggatttggg<br>agtttattattgcttccagagtaactatcttccgctcacgttcggtgctggcaccaagttggaaatcaaa |

| **Lineage c179 (IGHV1-42*01_IGKV4-80*01)** | |
|---|---|
| **Heavy chain sequence** | |
| germline | gaggtccagctgcagcagtctggacctgagctggtgaagcctggggcttcagtgaaga-<br>tatcctgcaaggcttctggttactcattcactggctactacatgaactgggtgaagcaaagtcctgaaaagagcct<br>tgagtggattggagagattaatcctagcactggtggtactacctacaac-<br>cagaagttcaaggccaaggccacattgactgtagacaaatcctccagcacagcctacatgcagctcaagagc<br>ctgacatctgaggactctgcagtctattactgtgcaagatccctctacggtagtag-<br>ttttcactggtacttcgatgtctggggcacagggaccacggtcaccgtctcctca |
| p7.BC76 | caggtgcagctgaaggagtctggacctgagctggtgaagcctggggcttcagtgaaga-<br>tatcctgcaaggcctctggttactcattcactgactattacatgaactgggtgaagcaaagtcctgaaaagagcc<br>ttgagtggattggagaggttaatcctaacactggtgatactacctataataa-<br>gaatttcaaggccaaggccacattgactgtagacaaatcctccaacacagcctacattcagctcaagagcctg<br>acatctgaggactctgcagtctattactgtgcaagatccctctacggtagtagtttccac-<br>tggtttttcgatgtctggggcacagggaccacggtcaccgtctcgagc |
| p7.BC49 | gaggtccagctgcaacagtctggacctgagctggtgaagcctggggcttcagtgaaga-<br>tatcctgcaaggcttctggttactcattcactgactactacatgaactgggtgaagcaaagtcctgaaaagagcc<br>ttgagtggattggagagattaatcctaacactggtgatactacctataataagag-<br>tttcaaggccaaggccacattgactgtagacaaatcctccaacacagcctacattcagctcaagagcctgaca<br>tctgaggactctgcagtctattactgtgcaagatccctctccggtagtagtttccac-<br>tggtttttcgatgtctggggcacagggactctggtcactgtctcgagc |
| p8.BC75 | gatgtgcagctgaaggagtctggacctgagctggtgaagcctggggcttcagtgaaga-<br>tatcctgcaaggcctctggttactcattcactgactattacatgaactgggtgaagcaaagtcctgaaaagagtct<br>tgagtggattggagaggttaatcctaacactggtgaaactacctataataa-<br>gaatttcaaggccaaggccacattgactgtagacaaatcctccaacacagcctacattcagctcaagagcctg<br>acatctgaggactctgcagtctattactgtgcaagatccctctacggtagtagtttccac-<br>tggtttttcgatgtctggggcacagggactctggtcactgtctcgagc |
| p8.BC46 | caggtgcagctgaagcagtctggacctgagctggtgaagcctggggcttcagtgaaga-<br>tatcctgcaaggcttctggttactcattcactgactactacatgaactgggtgaagcaaagtcctgaaaagagcc<br>ttgagtggattggagagattaatcctaacactggtgatactacctataataa-<br>gaatttcaaggccaaggccacattgactgtagacaaatcctccaacacagcctacattcagctcaagagcctg<br>acatctgaggactctgcagtctattactgtgcaagatccctctacggtagtagttttcac-<br>tggtttttcgatgtctggggcacagggactctggtcactgtctcgagc |
| **Light chain sequence** | |
| germline | caaattgttctcacccagtctccagcaatcatgtctgcatctctaggggagga-<br>gatcaccctaacctgcagtgccagctcgagtgtaagttacatgcactggtaccagcagaagtcaggcacttctc<br>ccaaactcttgatttatagcacatccaacctggcttctggagtcccttctcgcttcagtgg-<br>cagtgggtctgggaccttttattctctcacaatcagcagtgtggaggctgaagatgctgccgattattactgccatc<br>agtggagtagttatccatacacgttcggaggggggaccaagctggaaataaaa |

| | |
|---|---|
| p7.BC76 | gacatccagatgacacagtctccagcaatcatgtctgcatctctaggggagga-<br>gatcaccctaacctgcagtgccagttcgagtgtaaattacatgcactggtatcagcagaagtcaggctcttctcc<br>caaactcttgatttatagcacatccaacctggcttctggagtcccttctcgcttcagtgg-<br>cagtggatctgggaccttttattctctcacaatcagcagtgtggaggctgaagatgctgccgattattactgccatc<br>agtggagtagttatccatgtacgttcggagggggaccaagctggagctgaaa |
| p7.BC49 | gatatcgagatgacacagtctccagcaatcatgtctgcatctctaggggagga-<br>gatcaccctaacctgcagtgccagttcgagtgtaagttacatgcactggtatcagcagaagtcaggctcttctcc<br>caaactcttgatttatagcacatccaacctggcttctggagtcccttctcgcttcagtgg-<br>cagtggatctgggaccttttattctctcacaatcagcagtgtggaggctgaagatgctgccgattattactgccatc<br>agtggagtagttatccatgtacgttcggagggggaccagactggaaataaaa |
| p8.BC75 | caaattgtgatgacccagtctccagcaatcatgtctgcatctctaggggagga-<br>gatcaccctaacctgcagtgccagttcgagtgtaatttacatgcactggtatcagcagaagtcaggctcttctccc<br>aaactcttgatttatagcacatccaacctggcttctggagtcccttctcgcttcagtggcag-<br>tggatctgggaccttttattctctcacaatcagcagtgtggaggctgaagatgctgccgattattactgccatcagt<br>ggagtagttatccatgtacgttcggagggggaccagactggaaataaaa |
| p8.BC46 | caaattgttctcacccagtctccagcaatcatgtctgcatctctaggggagga-<br>gatcaccctaacctgcagtgccagttcgagtgtaagttacatgcactggtatcagcagaagtcaggctcttctcc<br>caaactcttgatttatagcacatccaacctggcttctggagtcccttctcgcttcagtgg-<br>cagtggatctgggaccttttattctctcacaatcagcagtgtggaggctgaagatgctgccgattattactgccatc<br>agtggagtagttatccatgtacgttcggaggggggtccaagctgaaaataaaa |

| **Lineage c87 (IGHV1-63*01_IGKV1-117*01)** | |
|---|---|
| **Heavy chain sequence** | |
| germline | caggtccagctgcagcagtctggagctgagctggtaaggcctgggacttcagtgaa-<br>gatgtcctgcaaggcttctggatacaccttcactaactactggataggttgggcaaagcagaggcctggacat<br>ggccttgagtggattggagatatttaccctggaggtggttatactaactacaatga-<br>gaagttcaagggcaaggccacactgactgcagacaaatcctccagcacagcctacatgcagttcagcagc<br>ctgacatctgaggactctgccatctattactgtgcaaggaaggtttactacggtagtac-<br>ctactttgactactggggccaaggcaccactctcacagtctcctca |
| p8.BC50 | caggtccagctgcagcagtctggagctgagctggtaaggcctgggacttcagtgaa-<br>gatgtcctgcaaggcttctggatacaccttcactaactactggataggttgggcaaagcagaggcctggacat<br>ggccttgagtggattggagatatttaccctggaaaaaattatactaactacaatgagaa-<br>gctcaagggcaaggccacactgacttcagacaaatcctccagcacagcctacatgcagttcagtagcctga<br>catctgaggactctgccatctattactgtgcaaggaaggtttactacggtagtacctacttt-<br>gactactggggccaaggcacccctctcacagtctcctca |
| p7.BC19 | caggtccaactccagcagcctggagctgacctggttaggcctgggacttcagtgaa-<br>gatgtcctgtaaggcttctggatacaccttcactaaccactggataggttggacaaagcagaggcctggacat<br>ggccttgagtggattggagatatttaccctggacgtgattatactaattacaatgagat-<br>tttcaagggcaaggccacactgactgcagacaaatcctccagcacagcctacatgcagttcagcagcctga<br>catctgacgactctgccatctattactgtgcaaggaaggtttactacggtagtacctacttt-<br>gactattggggccaaggaacctcagtcaccgtctcgagc |
| p8.BC83 | caggtccagctgcagcagtctggagctgacctggttaggcctgggacttcagtgaa-<br>gatgtcctgtaaggcttctggattcaccttcactaactactggataggttggacaaagcagaggcctggacatg<br>gccttgagtggattggagatatttaccctggacgtgattatactaattacaatgagat-<br>tttcaagggcaaggccacactgactgcagacaaatcctccagcacagcctacatgcagttcagcagcctga<br>catctgacgactctgccatctattactgtgcaaggaaggtttactacggtcatacctacttt-<br>gactactggggccaaggcaccactctcacagtctcctca |

| | |
|---|---|
| p7.BC27 | caggtgcaactgaagcagtctggagctgacctggttaggcctgggacttcagtgaa-gatgtcctgtaaggcttctggatacaccttcactaactactggataggttggacaaagcagaggcctggacatggccttgagtggattggagatatttaccctggacgtgattatactaattacaatga-gaatttcaagggcaaggccacactgactgcagacaaatcctccagcacagcctacatgcagttcagcagcctgacatctgacgactctgccatctattactgtgcaaggaaggtttactacggtagtac-ctactttgactactggggccaaggaacctcagtcactgtctcgagc |
| **Light chain sequence** | |
| germline | gatgttttgatgacccaaactccactctccctgcctgtcagtcttggagatcaa-gcctccatctcttgcagatctagtcagagcattgtacatagtaatggaaacacctatttagaatggtacctgcagaaaccaggccagtctccaaagctcctgatctacaaagtttccaaccgat-tttctggggtcccagacaggttcagtggcagtggatcagggacagatttcacactcaagatcagcagagtggaggctgaggatctgggagtttattactgctttcaaggttcacatgttccgtacacgttcg-gaggggggaccaagctggaaataaaa |
| p8.BC50 | gatgttttgatgacccaaattccactctccctgcctgtcagtcttggatatcaa-gcctccatctcttgcagatctagtcagagcattgtacatagtaatggaaacacctatttagaatggtatgtgcagaaaccaggccagtctccaaagctcctgatctacaaagtttccaaccgat-tttctggggtcccagacaggttcagtggcagcggatcagggacagatttcacactcaagatcagcagagtggaggctgaggatctgggagtttattactgctttcaaggttcacatgctccgtacacgttcg-gaggggggaccaacctggaaataaaa |
| p7.BC19 | gacattgtgatgacacagtctccactctccctgcctgtcagtcttggagatcaa-gcctccatctcttgcagatctagtcagagcattgtacatactaatggaaacacctatttagaatggtacctgcagaaaccaggccagtctccaaagctcctgatctacaaactttccaaccgat-tttctggggtcccagacaggttcagtggcagtggatcagggacagatttcacactcaagatcagcagagtggaggctgaggatctgggagtttattactgctttcaaggttcacatgttccgttcacgttcg-gaggggggaccagactggaaataaaa |
| p8.BC83 | gatgttttgatgacccaaactccactctccctgcctgtcagtcttggagatcac-gcctccatctcttgcagatctagtcagagcattgtacatagtaatggaaacacctatttagaatggtacctgcagaaaccaggccagtctccaaagctcctgatctacaaagtttccaaccgat-tttctggggtcccagacaggttcagtggcagtggatcagggacagatttcacactcaagatcagcagagtggaggctgaggatctgggagtttattactgctttcaaggttcacatgttccgttcacgttcg-gaggggggaccaagctggaaataaaa |
| p7.BC27 | agtgttgtgatgacccaaactccactctccctgcctgtcagtcttggagatcaa-gcctccatctcttgcagatctagtcagaacattgtacatagtaatggaaacacctatttagaatggtacctgcagaaaccaggccagtctccaaagctcctgatctacaaagtttccaaccgat-tttctggggtcccagacaggttcagtggcagtggatcagggacagatttcacactcaagatcagcagagtggaggctgaggatctgggagtttattactgctttcaaggttcacatgttccgttcacgttcg-gaggggggaccagactggaaataaaa |

**Supplementary Table 2 :**

| Antibodies | IGHV CDR mut.(aa) | IGHV FW mut.(aa) | IGKV CDR mut.(aa) | IGKV CFW mut.(aa) |
|---|---|---|---|---|
| germline.C179 | 0 | 0 | 0 | 0 |
| p7.BC76 | 5 | 7 | 2 | 5 |
| p7.BC49 | 5 | 5 | 1 | 5 |
| p8.BC75 | 5 | 8 | 2 | 3 |
| p8.BC46 | 4 | 7 | 1 | 3 |
| germline.C127 | 0 | 0 | 0 | 0 |
| p9.BC41 | 1 | 4 | 0 | 4 |
| p9.BC18 | 0 | 2 | 0 | 1 |
| p10.BC50 | 1 | 2 | 0 | 3 |
| germline.c87 | 0 | 0 | 0 | 0 |
| c87.p8.BC50 | 2 | 3 | 1 | 4 |
| c87.p7.BC19 | 3 | 7 | 0 | 7 |
| C87.p8.BC83 | 4 | 4 | 0 | 3 |
| C87.p7.BC27 | 2 | 8 | 1 | 4 |

### SUPPLEMENTARY FIGURES

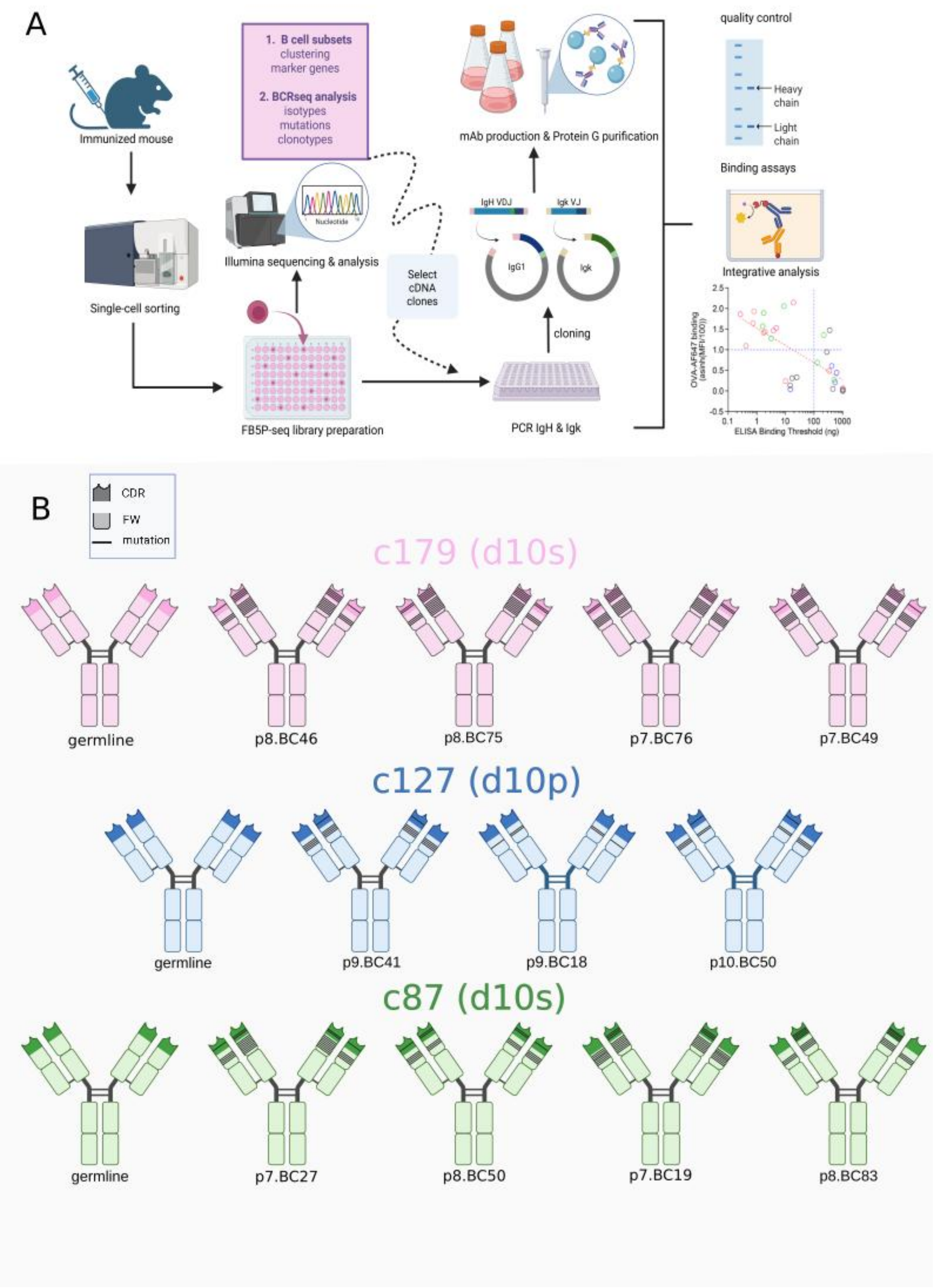


**Supplementary Figure 1 : Immunization, production and sequences of three antibody lineages**

**(A)** Antibody production workflow. **(B)** Schematic representation of the three antibody lineages of this study, indicating the time points (days post-immunization) at which the corresponding lymph nodes were collected. Each lineage consists of a germline and its mutated antibodies. Each black stripe represents a mutation acquired during maturation; its presence over a light color indicates belonging to a FW part of the antibody; its presence over a dark color indicates belonging to a CDR part of the antibody.

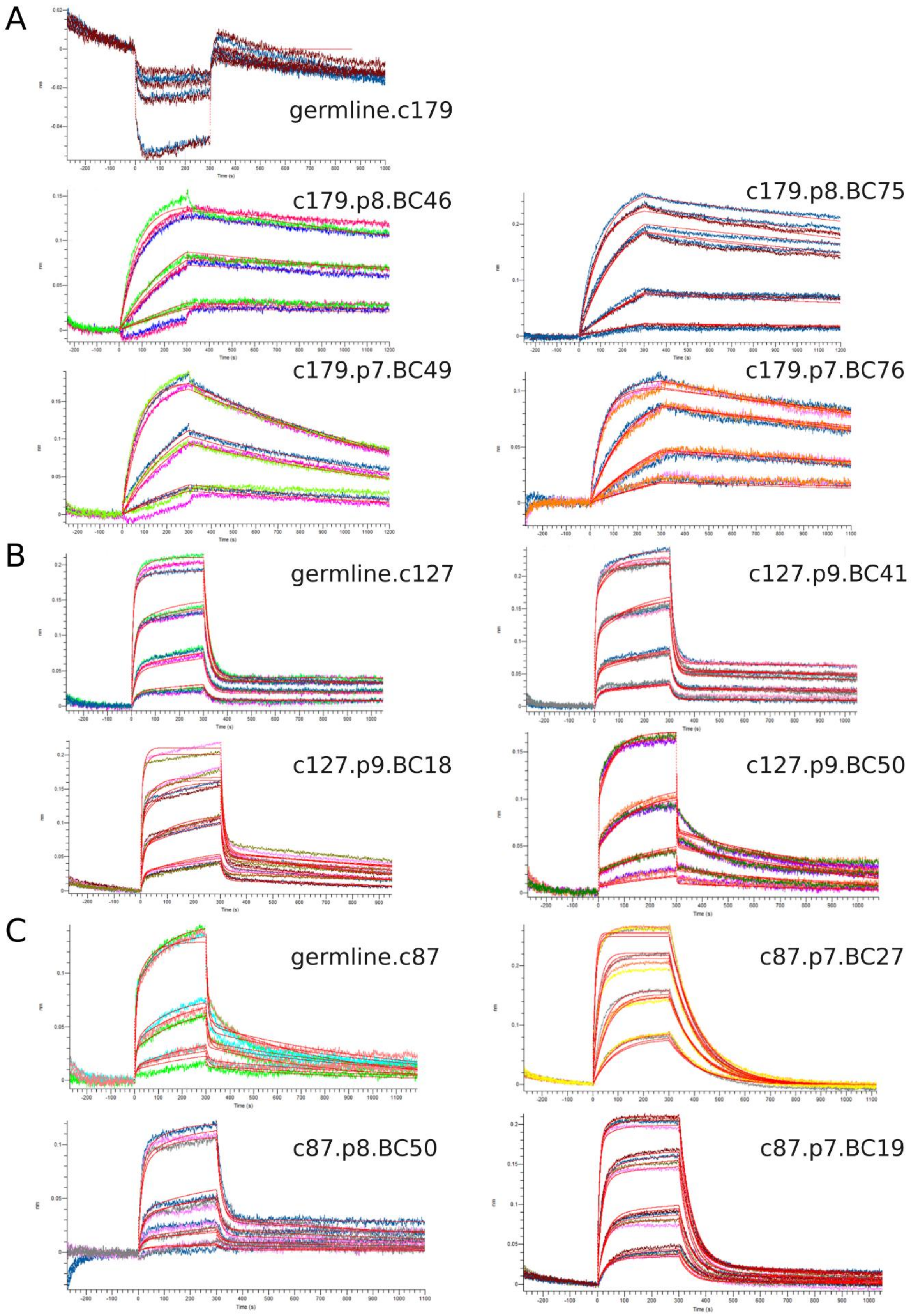


**Supplementary Figure 2 : Sensorgrams of binding in solution obtained by BioLayer Interferometry of every antibody measured. (A)** Schematic of the biolayer interferometry assay. **(B)** Recorded binding curves (sensorgrams) from the association and dissociation phases were analyzed to extract kinetic parameters. Sensorgrams represent the shift in wavelength of the spectrum reflected by the sensor tip versus time, with association starting at the moment of contact of tip with ligand solution and resulting in an increasing signal, and dissociation starting at the following contact of tip with solvent solution alone and resulting in a decreasing signal. Experiments were conducted over three or four concentrations that span the binding regime (the level of signal increasing with the concentrations) forming a total of three or four sets of superimposing curves per graph. In each graphic, replicate experiments at a given concentration are denoted by traces with different colors forming a set of broadly superimposing curves. A minimum of three replicate experiments was performed per antibody. Each kinetic responses were fitted to the 1:1 Langmuir kinetic model to obtain values for association ($k_{on}$), dissociation ($k_{off}$), and the equilibrium dissociation constant ($k_d$). The globally fitted curves are represented in red lines. **(C)** represents the sensorgrams from the association and dissociation phases of c179 lineage and its germline antibody. For each antibody, C179 germline makes an exception, as no signal was obtained, resulting in indistinguishable curves. **(D)** shows similar types of data for all antibodies of the c127 lineage. **(E)** shows similar types of data for all antibodies of the c87 lineage.

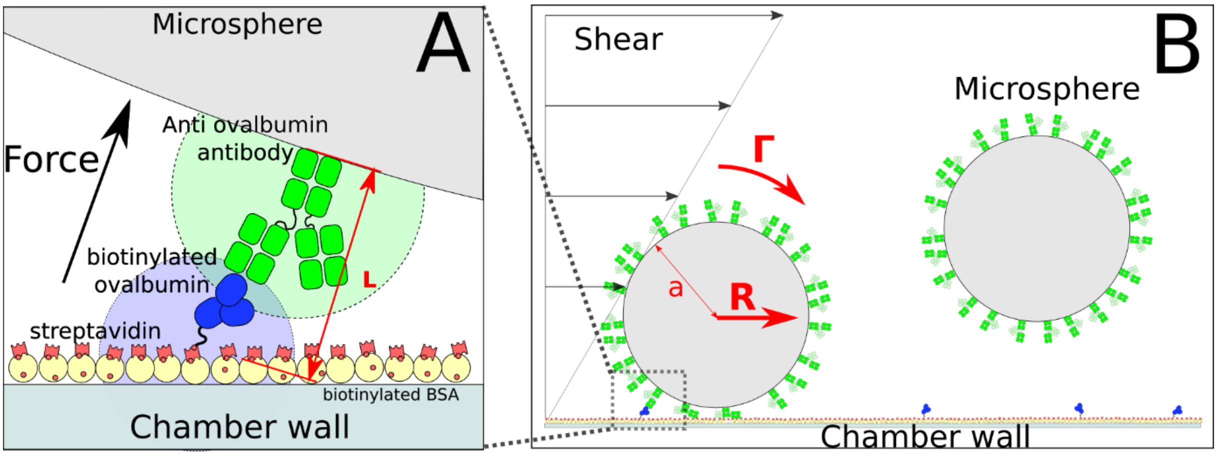


**Supplementary Figure 3 : Laminar flow chamber principle and apparatus (A)** Principle of the laminar flow chamber: Microspheres are decorated with a high amount of antibody; chamber surface is decorated with low amounts of ovalbumin. A single antibody-ovalbumin bond may stop the microsphere in the range of forces applied by shear force on it. **(B)** Force applied on the bond depended on shear flow that produced both a hydrodynamic drag *R*, and a torque *Γ* on the microsphere, depending on medium viscosity and microsphere radius a. A lever effect makes the actual force F applied on the bond dependent on the length of the molecular bond L. Force on bond was calculated as $F = \left(R + \frac{\Gamma}{a}\right) \times \sqrt{\frac{a}{2l}}$ with $R = 1.7005 \times 6\pi\mu a^2 G$ and $\Gamma = 0.9440 \times \pi\mu a^3 G$ (with $R$ the traction on the microsphere, $\Gamma$ the torque on the microsphere, $a$ the microsphere radius, $l$ the total bond length (24nm), µ the medium viscosity (7x10-4 Pa.s at 37°C), and $G$ the shear rate.

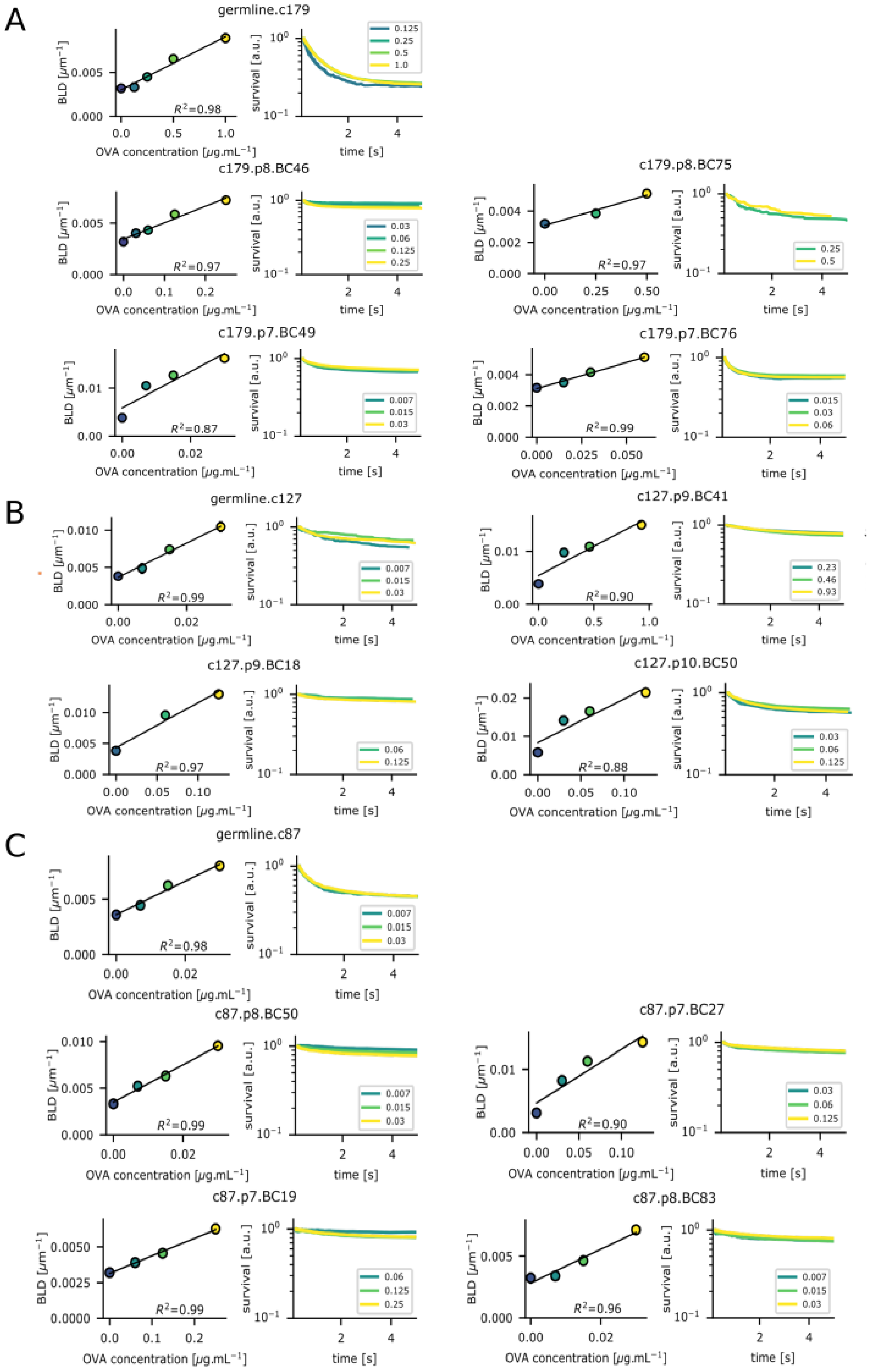


**Supplementary Figure 4 : Assessment of single bond observation for all the antibodies measured. (A)** For each antibody of the c179 lineage and its germline antibody is a couple of two plots: The scatter plot on the left shows interpolated BLD values as a function of the concentration of ovalbumin at a shear rate corresponding to a 10pN force on the bond. Errors bars show standard error. BLD is proportional to the amount of deposited ovalbumin in the single molecular bond range (Spearman $R^2$ determination coefficient is shown.) The right panel shows the logarithm of the fraction of surviving bonds plotted against the duration of the bonds under a force of 10pN. In the range of single molecular binding, survival is independent of the amount of deposited ovalbumin and corresponding curves thus superimpose. **(B)** Similar types of data for all antibodies of the c127 lineage. **(C)** Similar types of data for all antibodies of the c87 lineage.

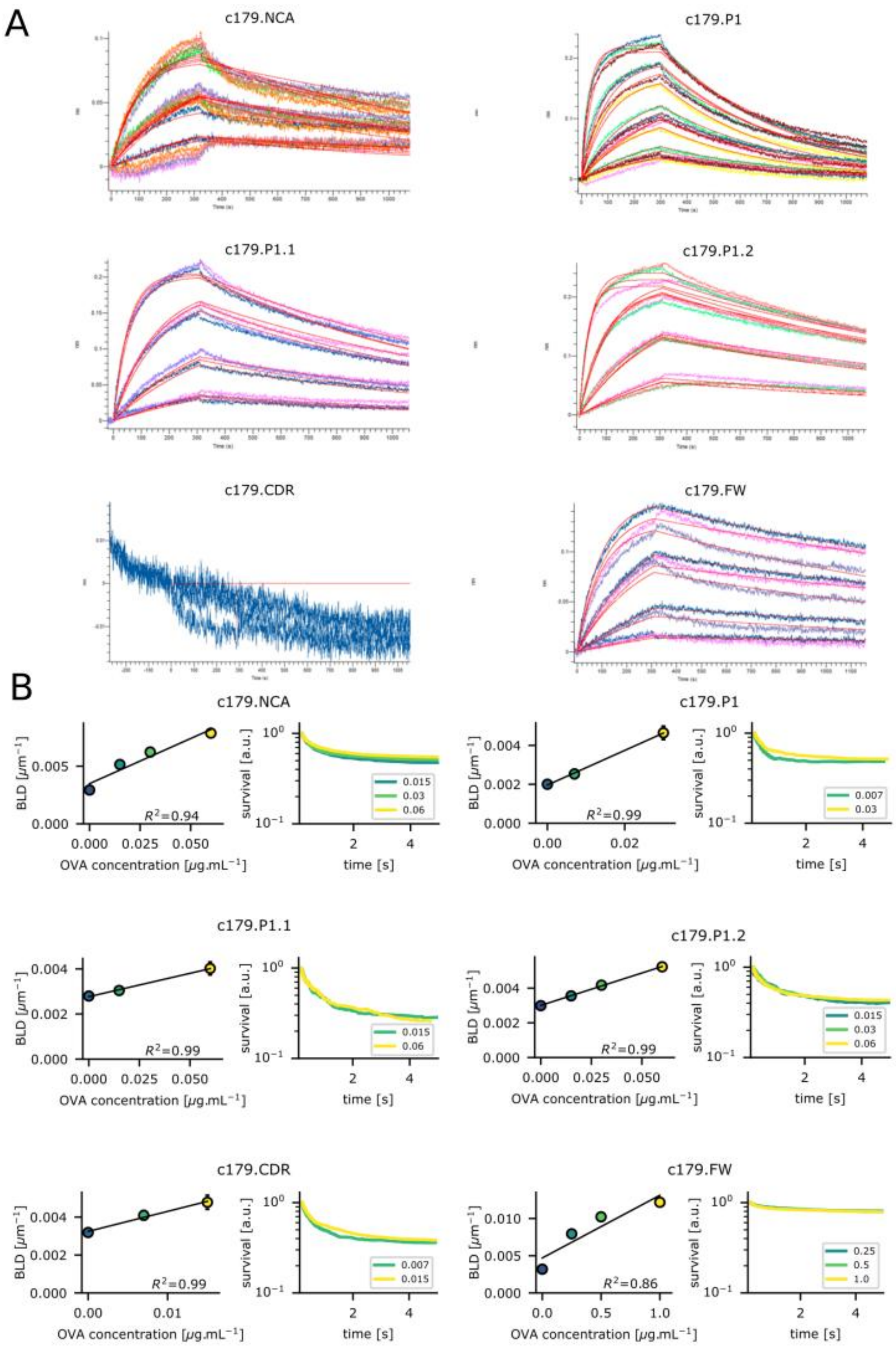


**Supplementary Figure 1: Sensorgrams of binding in solution obtained by BioLayer Interferometry and single bond observations data by Laminar Flow Chamber for c179 reverted antibodies. (A)** represents the sensorgrams from the association and dissociation of data for all antibodies of c179

lineage with partially reverted maturation-acquired mutations. C179 CDR-/FW+ makes an exception, as no signal was obtained, resulting in indistinguishable curves. **(B)** For the same set of antibodies (c179 lineage with partially reverted maturation acquired mutations antibodies) represents on the left the scatter plots that show the interpolated BLD values as a function of concentration of ovalbumin at a shear rate corresponding to a 10pN force on the bond. Errors bars show standard error. BLD is proportional to the amount of deposited ovalbumin in the single molecular bond range (Spearman $R^2$ determination coefficient is shown). The right panel shows the logarithm of the fraction of surviving bonds plotted against the duration of the bonds under a force of 10pN. In the range of single molecular binding, survival is independent of the amount of deposited ovalbumin and corresponding curves thus superimpose.
